# Unveiling the functional heterogeneity of endogenous tissue extracellular vesicles in skeletal muscle through multi-omics

**DOI:** 10.1101/2024.11.20.624461

**Authors:** Yizhuo Wang, Peng Lou, Xiyue Zhou, Yijing Xie, Yimeng Zhang, Shuyun Liu, Lan Li, Yanrong Lu, Meihua Wan, Jingqiu Cheng, Jingping Liu

**Author notes:** Co-first authors that contributed equally to this work (Yizhuo Wang, Peng Lou). Corresponding Author: Jingping Liu Jingqiu Cheng Address: NHC Key Laboratory of Transplant Engineering and Immunology, West China Hospital, Sichuan University, No. 2222 Xinchuan Road, Chengdu 610041, China.

## Abstract

Extracellular vesicles (EVs) have emerged as promising tools for the development of disease biomarkers and therapeutics because they can transfer various bioactive cargos between cells *in vivo*. A better understanding of the heterogeneous properties of EVs *in vivo* may provide insights into their biological roles and clinical translation potential. As a proof-of-concept, we report that different EV subpopulations from skeletal muscle tissues have distinct composition signatures and diverse biological effects on recipient cells. Multiple cell types (e.g., myoblasts and endothelial cells (ECs)) can contribute to the pool of muscle tissue-derived EVs, and large EVs (L-EVs) are enriched with proteins related to metabolic regulation, whereas small EVs (S-EVs) are enriched with original muscle cell-specific proteins related to muscle function regulation. Compared with L-EVs, S-EVs exhibited abundant surface proteins and higher cell uptake rates. Moreover, L-EVs and S-EVs can induce diverse changes in global gene expression, metabolic patterns and some cellular behaviors (e.g., proliferation and differentiation) in recipient cells. These results suggest that different EV subpopulations might control tissue hemostasis in a coordinated manner and suggest the importance of reconsidering their favorable role in future applications (e.g., S-EVs for biomarker discovery and L-EVs for metabolic intervention). This study highlights the functional heterogeneity of tissue-derived EVs *in vivo*, and the selection of an ideal EV subset on the basis of its specific biological properties may be a promising strategy for developing more precise biomarkers or tailored EV therapies for regenerative medicine.

**Graphic abstract:** 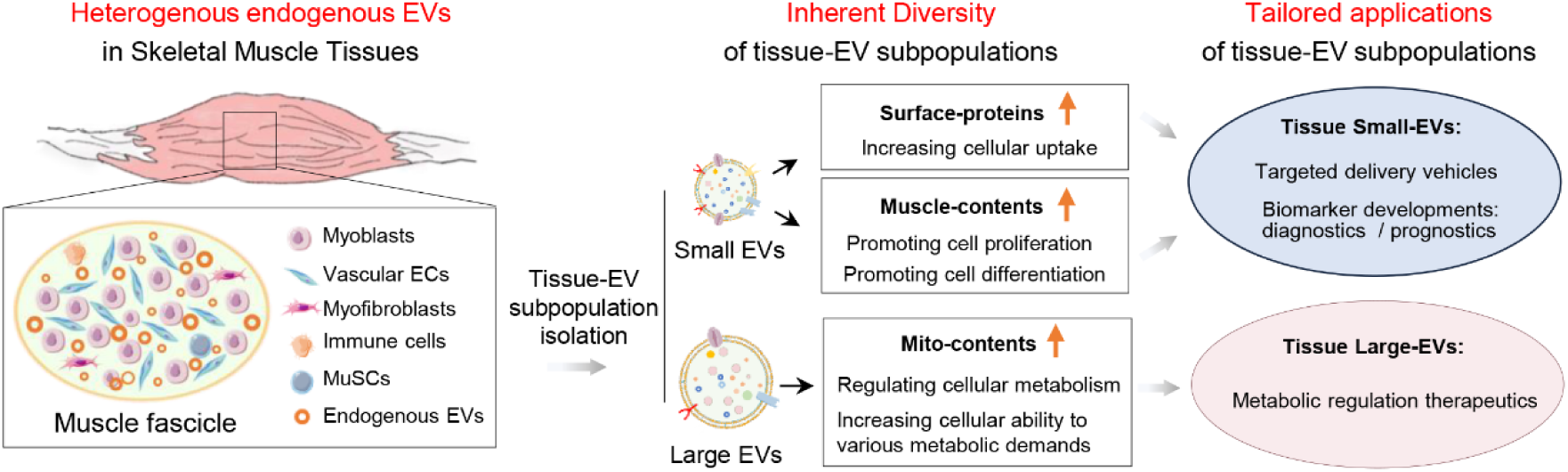

**Significance:** Extracellular vesicles (EVs) are emerging as important tools for diagnostics, therapeutics, and prognostics in various diseases. Understanding the inherent heterogeneity of EVs is crucial, as distinct subpopulations function differently. While extensive research focuses on EVs derived from cell supernatants, endogenous tissue EVs can more accurately reflect the pathophysiological characteristics of their originating cells. Here, we propose different tissue-EV subpopulations coordinately regulate tissue homeostasis. Small EVs with tissue-specific signatures show promise for biomarker development, while large EVs with metabolic signatures are suitable for metabolic interventions. Additionally, small EVs with enhanced surface proteins are ideal for targeted delivery. This work highlights the importance of selecting tissue-EV subpopulations based on their unique properties for developing precise biomarkers and tailored therapies in regenerative medicine.

## Introduction

Notably, extracellular vesicle (EV) secretion is a highly conserved biological process in nearly all cells. Since EVs mediate cell‒cell or tissue‒tissue communication in various biological processes (e.g., tissue homeostasis controlling, immune responses, and metabolic regulation), they have been considered as promising tools for disease biomarker discovery or therapeutic development. Traditionally, EVs have been categorized into certain subtypes (e.g., exosomes, microvesicles, and apoptotic bodies) on the basis of their biogenesis routes [1, 2]. Owing to the heterogeneous nature of EV biology, different EV subpopulations have varied sizes, compositions, and biological functions. For example, mesenchymal stem cell (MSC)-derived exosomes and microvesicles have distinct transcriptome profiles [3]. However, the existing findings concerning EV heterogeneity are derived mainly from *in vitro* cultured cell-derived EVs (cell-EVs). Because metabolic states and EV secretion can be readily affected by culture conditions, cell-EVs may have altered properties and cannot accurately reflect the status of their original tissue cells *in vivo* [4, 5]. In fact, body tissues harbor large numbers of EVs secreted by resident cells and infiltrating cells [4, 5], and these tissue cell-derived endogenous EVs (tissue-EVs) may act as versatile signaling conduits to regulate tissue remodeling and injury repair processes[4, 6]. Therefore, an in-depth understanding of the heterogeneous nature of tissue-EVs may provide insights into the regulatory mechanism of tissue homeostasis and repair, as well as the development of EV-based biomarkers or therapies for precise regenerative medicine.

To reveal the functional heterogeneity of tissue-EVs, skeletal muscle was selected as a target tissue in the current study, since it is one of the largest body tissues (∼40% of the total body weight) and serves as a major protein reservoir [7]. Moreover, skeletal muscle can dynamically remodel in response to physiological or pathological stimuli and secrete many soluble factors and EVs to regulate systemic metabolic states, locomotion, thermoregulation, etc. Recent studies have indicated that tissue-EVs play vital roles in muscle-mediated local and systemic tissue homeostasis and that ∼5% of circulating blood EVs might originate from skeletal muscle because they contain skeletal muscle-specific cargos [8]. The levels of circulating EVs markedly increase after aerobic exercise, suggesting the dynamic biological roles of tissue-EVs in skeletal muscle under different conditions [9]. Furthermore, skeletal muscle cell-derived EVs can interact with other cell types. For example, myogenic progenitor cell-derived EVs were shown to reduce extracellular matrix (ECM) production in interstitial fibrogenic cells during muscle hypertrophy[10]. Together, these findings suggest that skeletal muscle-derived tissue-EVs play vital roles in adapting to stimuli and regulating tissue homeostasis both locally and systemically, making them an ideal model for exploring the functional heterogeneity of tissue-EV subpopulations.

Here, we report that different muscle tissue-EV subpopulations have distinct composition signatures and can induce diverse biological responses in recipient cells. Healthy skeletal muscle tissue-derived large EVs (L-EVs) were enriched with proteins involved in metabolic regulation, whereas small EVs (S-EVs) were enriched with muscle cell-specific proteins involved in muscle function regulation. Compared with L-EVs, S-EVs had abundant surface proteins and higher cell uptake rates. Moreover, L-EV and S-EV treatments could induce diverse changes in global gene expression, metabolic patterns and cell behavior in recipient cells, and they might control muscle tissue hemostasis in a coordinated manner. This study highlights the functional heterogeneity of tissue-EVs and suggests that the selection of a suitable subpopulation with specific biological properties may be a promising strategy for developing more precise biomarkers or tailored EV therapies.

## Results and discussions

### L-EVs and S-EVs from healthy muscles have distinct composition signatures

To date, EVs have shown potential in disease diagnostic or therapeutic applications, because they carry various original cell-derived bioactive cargos[11]. Body tissues serve as native reservoirs of endogenous EVs released by diverse cell types, and these tissue-EVs can act on proximal cells or cruise through the circulatory system to impact distal cells *in vivo* by transferring functional cargos [5]. However, the detailed compositions and biological roles of different subpopulations of tissue-EVs are incompletely understood. Although some previous studies termed their prepared EVs as exosomes or microvesicles, it is impossible to assign the isolated EVs to a particular biogenesis pathway after collection [1]. According to the Minimal Information Guidelines for Extracellular Vesicle Studies (MISEV2023), terminologies, such as “exosomes” and “microvesicles”, which are based on biogenesis routes to define isolated EVs, may be misleading and are difficult to establish [1]. Instead, terms based on size-cut-offs, such as “small EVs” and “large EVs”, have been recommended and widely used in recent studies despite some overlap may exist[1, 12]. In line with these reports, TEM images of healthy mouse skeletal muscle revealed the presence of numerous tissue-EVs of various sizes in the interstitial spaces among cells (Figure 1A-B). Both L-EVs (size > 150 nm) and S-EVs (size < 100 nm) could be observed within such muscle tissues (Figure 1B). Next, L-EVs and S-EVs were isolated from healthy mouse skeletal muscle tissues via an optimized differential ultracentrifugation method with slight modifications [6] (Figure 1A). The TEM results revealed that both EV subtypes were typical bilayer vesicles and that the median size of S-EVs (∼95 nm) was smaller than that of L-EVs (∼151 nm) (Figure 1C-1D). Despite a few overlaps in the size distribution between 100∼150 nm, ∼52.6% of the S-EVs were smaller than 100 nm, whereas ∼51.2% of the L-EVs were larger than 150 nm (Figure 1E). These results indicate the successful isolation of EV subpopulations from skeletal muscles, which aligns with the recommendations of MISEV2023.

**Figure 1.**
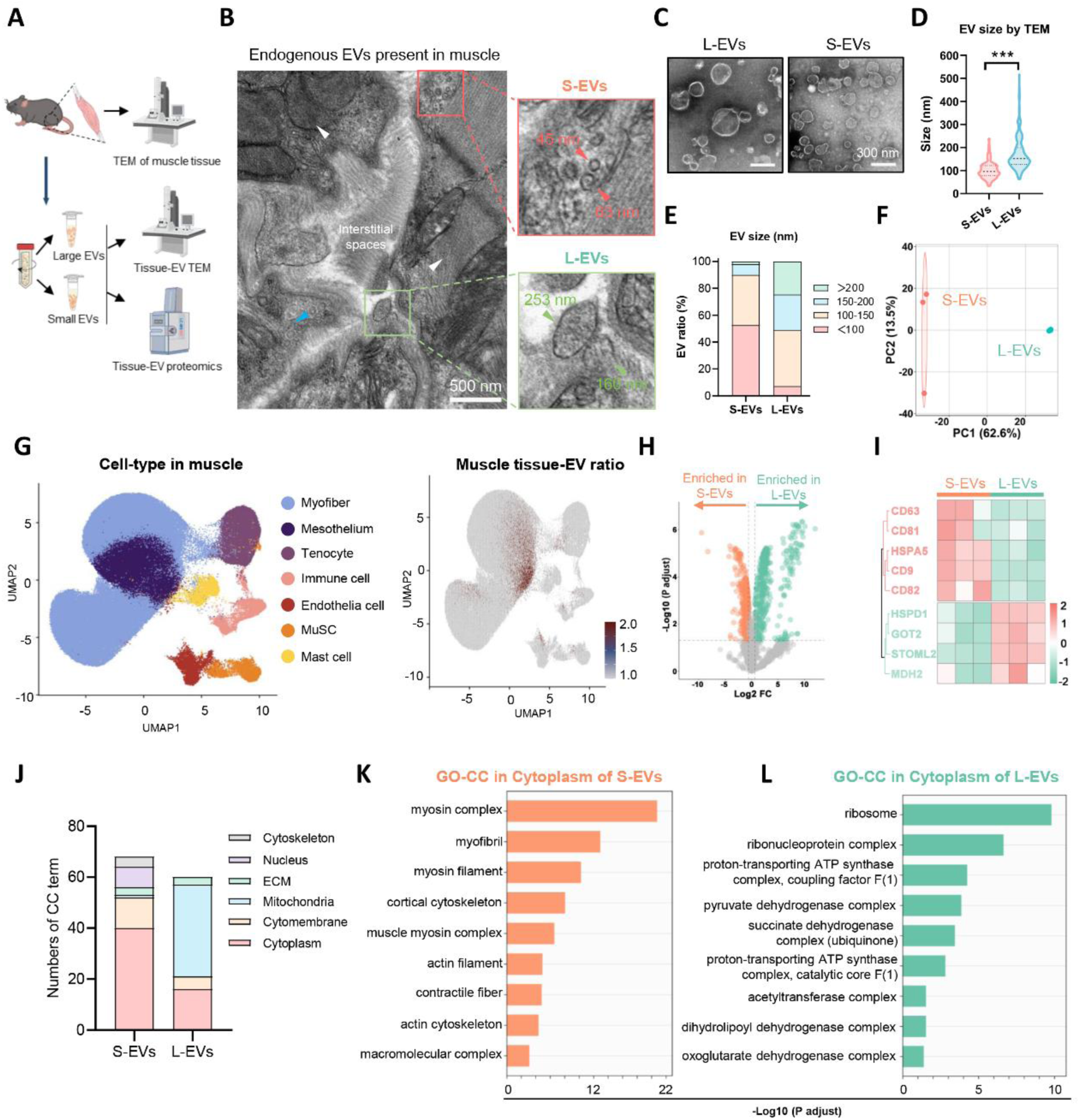
Isolation and characterization of skeletal muscle tissue-EV subsets. (A) Schematic illustration of the muscle tissue-EV heterogeneity investigation. (B) Representative TEM image of endogenous EVs within mouse skeletal muscle tissues (the red box indicates S-Evs, the green box indicates L-Evs, the white arrows indicate muscle cells, and the blue arrows indicate Ecs; scale bar = 500 nm). (C) Representative TEM image of L-EV and S-EV preparations from skeletal muscle tissues (scale bar = 300 nm). (D-E) EV size distribution determined based on TEM images. (F) PCA scatter plot of different groups based on proteomic data showing the differences between different groups (n = 3). (G) Mapping detection of cell type-specific proteins in clusters of tissue Evs. Uniform manifold approximation and projection (UMAP) visualization of annotated cells in the Muscle Cell Atlas (left panel) and EV proteomic mapping with the muscle cell atlas (right panel). (H) Volcano plots showing the DEPs (FC > 1.2 and p-adjusted < 0.05) between L-EVs and S-EVs (n = 3). (I) Heatmap of selected proteins enriched in L-EVs or S-EVs from the muscle tissues. (J) Sublocalization classification of GO-CC analysis of DEPs enriched in S-EVs or L-EVs. (K) GO-CC analysis of cytoplasmic terms enriched in S-EVs. (L) GO-CC analysis of cytoplasmic terms enriched in L-EVs.

Preliminary studies have revealed that different subpopulations of cell-EVs might have diverse compositions. For example, cancer cell (pleural mesothelioma cell)-derived L-EVs (pelleted at 10 K ×g) were enriched with mitochondrial components, whereas S-EVs (pelleted at 100 K ×g) were enriched with endoplasmic reticulum (ER)-related components [12]. Due to the high complexity of tissue cell types and their microenvironments [13], different subpopulations of tissue-EVs may have distinct cargos related to diverse biological effects. The global protein compositions of L-EVs and S-EVs isolated from healthy mouse muscle were profiled via liquid chromatography–coupled tandem mass spectrometry (LC‒MS/MS)-based proteomics. A total of 2128 proteins were identified in the tissue-EVs, and L-EVs and S-EVs clearly separated and exhibited an overall differential protein expression pattern (Figure 1F, Figure S1). To explore the cell origins of such EVs, we mapped the proteomic data of tissue-EVs with the public single-cell atlas of mouse skeletal muscle as previously reported [14], and the main proportions of tissue-EVs were likely from myoblasts and ECs (two of the major resident cells in skeletal muscles), followed by immune cells, tenocytes, and MuSCs (Figure 1G), suggesting that multiple cell types contribute to the pool of muscle tissue-EVs. The differentially expressed proteins (DEPs; *P*-adjusted < 0.05) between L-EVs and S-EVs, including 466 DEPs with higher expression in L-EVs (vs. S-EVs) and 435 DEPs with higher expression in S-EVs (vs. L-EVs), were identified (Figure 1H). A previous study revealed that cultured breast cancer cell-derived S-EVs contain abundant tetraspanins (e.g., CD63, CD81, and CD9) and ESCRT (endosomal sorting complex required for transport) proteins, whereas L-EVs contain abundant mitochondrion- and cytokinesis-associated proteins (PRC1, TIM/TOM) [15]. Similarly, S-EVs from healthy muscle tissues contained abundant tetraspanins (e.g., CD63, CD81, CD9, and CD82), whereas L-EVs contained abundant metabolic proteins (e.g., HSPD1, GOT2, STOML2, and MDH2) (Figure 1I). These results indicate that different tissue-EV subpopulations from the same cell origin have distinct compositions.

The subcellular origins of the EVs were evaluated via gene ontology (GO) cellular component (CC) analysis. Compared with L-EVs, S-EVs presented higher levels of cytoplasm, cytomembrane, and nucleus proteins, whereas L-EVs presented higher levels of mitochondria, cytoplasm, and ECM proteins than S-EVs did (Figure 1J). The The expression pattern of cytoplasmic proteins (the largest CC fraction) between S-EVs and L-EVs was further assessed, and S-EVs carried abundant cytoplasmic proteins related to cytoskeletal and muscle functions, such as the myosin complex, myofibril, myosin filament, and cortical cytoskeleton (Figure 1K), whereas L-EVs carried abundant cytoplasmic proteins linked to metabolic pathways and protein synthesis, such as the ribosome, proton-transporting ATP synthase complex, pyruvate dehydrogenase complex, and succinate dehydrogenase complex. (Figure 1L). Together, these results indicate that L-EVs and S-EVs have different composition signatures and subcellular origins, which may be associated with diverse biological functions in recipient cells.

### L-EVs are enriched with mitochondrial proteins engaged in metabolic regulation

Next, we explored the composition signature and possible biological role of L-EVs compared with those of S-EVs. KEGG pathway enrichment analysis revealed that the DEPs of L-EVs (vs. S-EVs) were involved in many metabolic pathways, such as oxidative phosphorylation, carbon metabolism, the citrate (TCA) cycle, and reactive oxygen species (Figure 2A). Mitochondria are central organelles that control cellular metabolic and redox pathways [16, 17]. The mitochondrial proteins in tissue-EVs were further assayed using mitoCarta2.0 [18], and ∼25.64% (529 of 2063) of the EV proteins were mitochondrial proteins (Figure 2B). Among these proteins, ∼43.37% (229 of 529) were from the mitochondrial inner membrane (MIM), ∼38.07% (201 of 529) were from the mitochondrial matrix, and ∼8.71% (46 of 529) were from the mitochondrial outer membrane (MOM) (Figure 2B). Compared with S-EVs, L-EVs presented an overall increase in the levels of mitochondrial proteins (MOM, matrix, MIM, intermembrane space (IMS), and membrane) (Figure 2C). MIM participates in coordinating cellular energetics and apoptotic pathways by integrating the electron transport chain and ATP synthesis machinery, and the mitochondrial matrix is the major site of the TCA cycle and fatty acid oxidation [19]. The enriched MIM proteins of L-EVs (vs. S-EVs) were involved in energy metabolism pathways, such as mitochondrial ion regulation and oxidative stress response, mitochondrial structure and function maintenance (Figure 2D-E); the matrix proteins of L-EVs were involved in nutrients catabolism and energy metabolism pathways (Figure 2F-G); the MOM proteins of L-EVs were involved in mitochondrial organization, anion/protein transport, apoptotic process, oxidative process (Figure 2H-J). Collectively, these results suggest that L-EVs are enriched with mitochondrial proteins, which might be associated with metabolic regulation processes.

**Figure 2.**
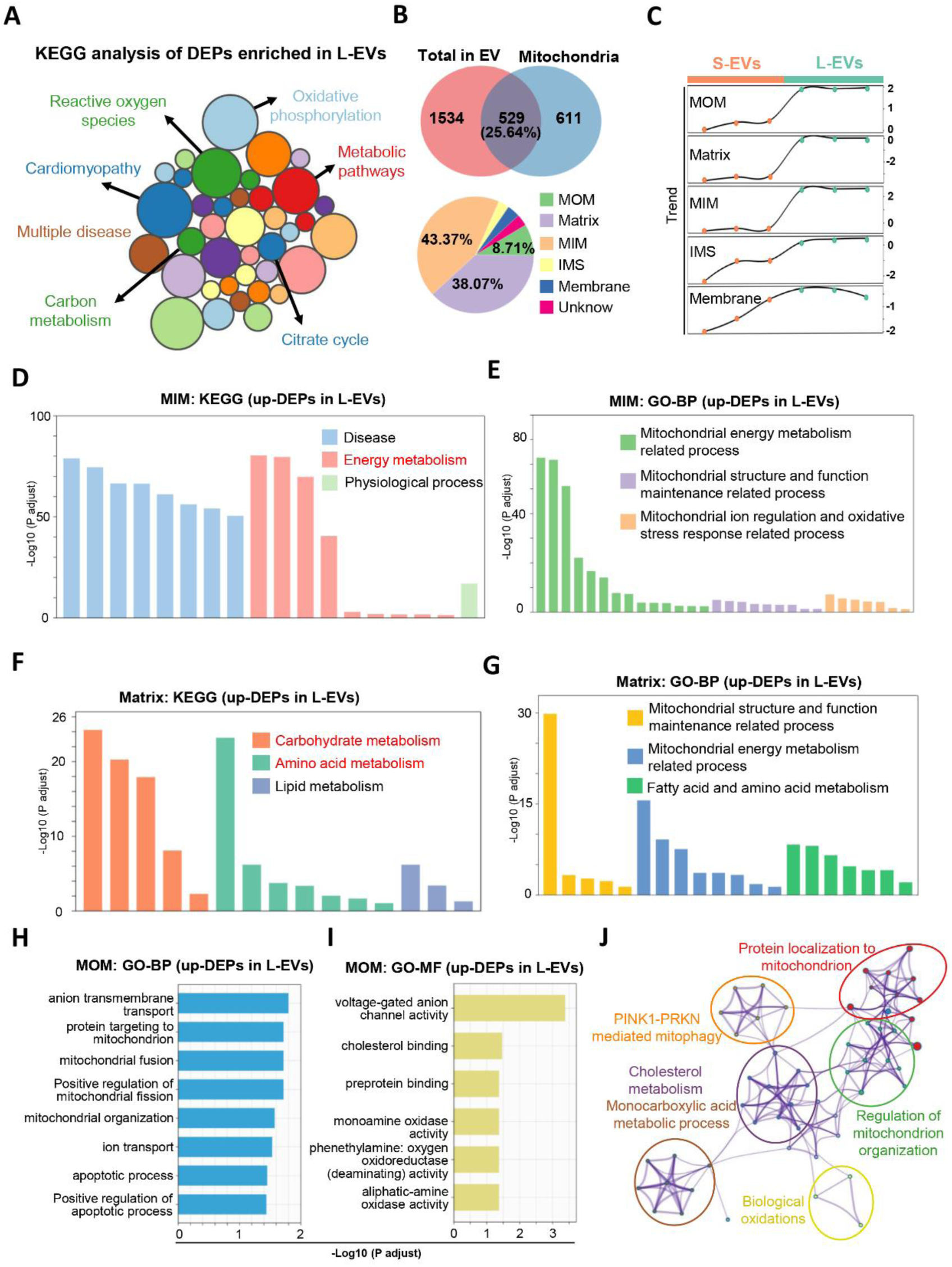
Evaluating the composition signatures and biological functions of L-EVs. (A) KEGG enrichment analysis of the upregulated DEPs in L-EVs compared to S-EVs. (B) Pie charts depicting the number (top) and sublocalization (bottom) of mitochondrial DEPs (mito-DEPs) revealed by proteomic analysis in tissue-EVs (n = 3). (C) Trend analysis of protein expression levels in each sublocalization region between L-EVs and S-EVs. (D-E) KEGG enrichment GO biological process analysis of the DEPs of L-EVs (vs. S-EVs) localized in the MIM. (F-G) KEGG enrichment and GO biological process analysis of the DEPs of L-EVs (vs. S-EVs) localized in the matrix. (H-J) GO biological process analysis and interaction network analysis of the DEPs of L-EVs (vs. S-EVs) localized in the MOM.

In recent years, an increasing number of studies have shown that EVs can carry functional mitochondrial contents (e.g., mitochondrial proteins, mtDNA) and transfer them into recipient cells to impact the metabolic states or phenotypes of these cells [16, 20]. For example, serum EVs enriched with mtDNA could enhance mitochondrial oxidative phosphorylation (OXPHOS), increase mtROS levels, and increase the mitochondrial membrane potential in epithelial cells [21]. On the basis of these reports, healthy-tissue L-EVs might function primarily in maintaining cellular energy and redox homeostasis, regulating cellular adaptation under physiological stimuli by supporting metabolic plasticity. EV-mediated mitochondrial content release is a potent mechanism of mitochondrial quality control that aims to remove injured or disabled mitochondria from donor cells [22]. In this context, the release of muscle mitochondria-rich L-EVs may be an instinct response to adapt to cellular metabolic challenges and oxidative stress. However, this concept cannot fully address the fact that healthy cells or tissues under steady-state conditions can also secrete large amounts of mitochondria-rich EVs. Additionally, it is still unclear why muscle tissue cell-derived mitochondrial proteins are primarily sorted into L-EVs rather than S-EVs. Thus, future studies are needed to explore the biogenesis routes of L-EVs and their detailed role in the metabolic regulation of target cells.

### S-EVs are enriched with muscle-specific proteins engaged in muscle regulation

The distinct compositions and biological roles of S-EVs compared with L-EVs were also explored. GO analysis revealed that the upregulated proteins in S-EVs (vs. L-EVs) were enriched in components related to muscle (∼34.69%), the cytoskeleton (∼22.45%), and the endoplasmic reticulum (ER, ∼12.24%) (Figure 3A-B). Specifically, S-EVs presented higher levels of proteins related to muscle cells than L-EVs did (Figure 1J). Similarly, adipocyte-derived S-EVs presented higher levels of adipocyte-specific proteins (e.g., adiponectin) than L-EVs did [23]. We further explored whether tissue EVs express the identified markers of the major tissue cell types within skeletal muscle as previously reported [7], and whether S-EVs expressed greater numbers of the identified markers of myoblasts or ECs than L-EVs did. The findings suggest that, compared with L-EVs, S-EVs may carry more original tissue cell-specific cargos and exhibit greater similarity (Figure S2A-S2C). The possible biological role of S-EV proteins was further evaluated via two major component terms (muscle and cytoskeleton). GO enrichment analysis revealed that S-EVs were involved mainly in biological processes related to muscle contraction and muscle development, cell motility and cytoskeleton organization, and DNA repair and remodeling (Figure 3C and 3E). KEGG enrichment analysis revealed that S-EVs might participate in pathways related to the regulation of the actin cytoskeleton, motor protein, focal adhesion, biosynthesis of glycosylphosphatidylinositol (GPI)-anchor and N-glycan, protein processing in the ER, etc. (Figure 3D and 3F).

**Figure 3.**
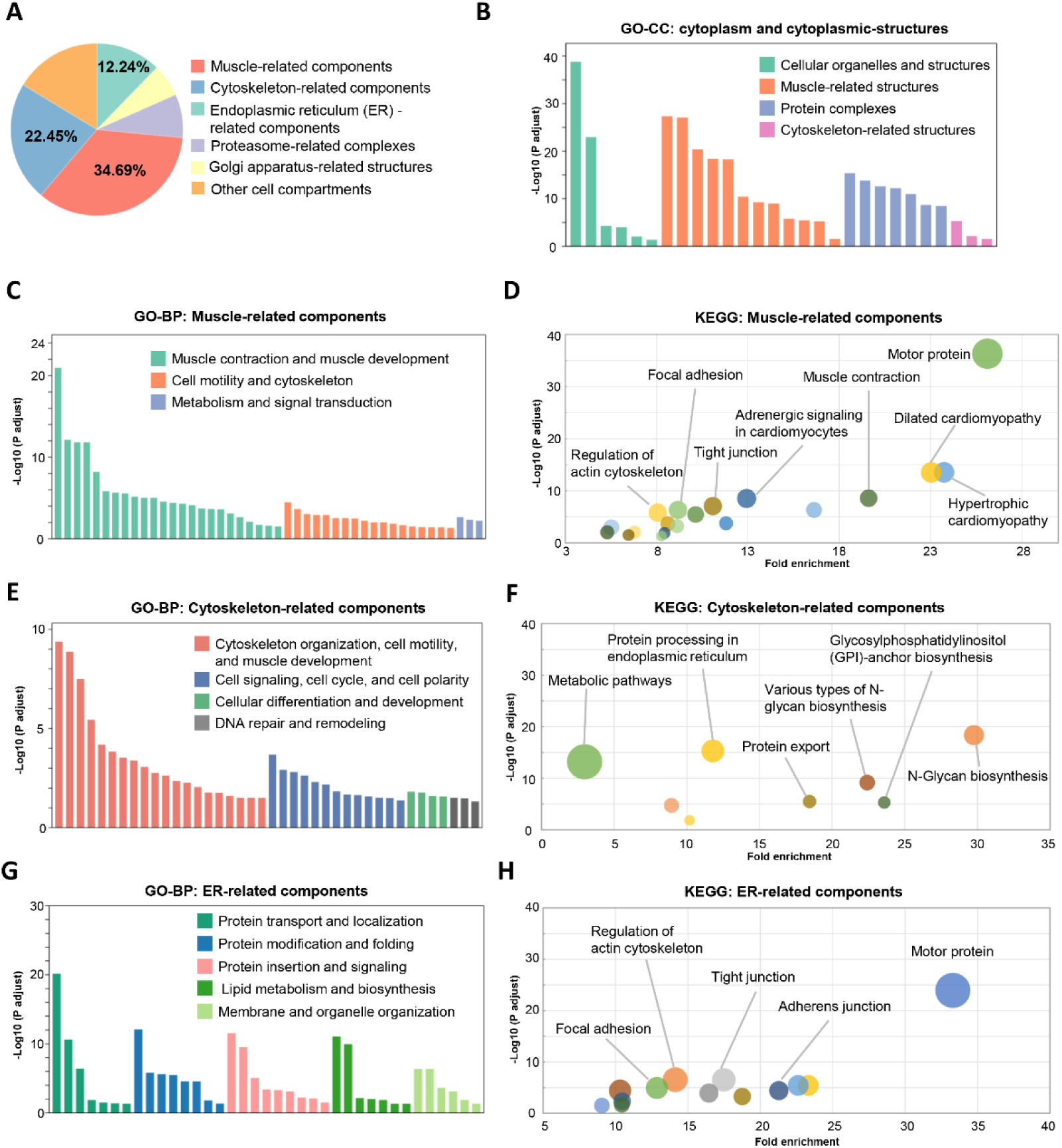
Evaluating the composition signatures and biological functions of S-EVs. (A) Pie chart depicting the sublocalization of DEPs among the upregulated DEPs in S-EVs (vs. L-EVs) by GO cellular component analysis. (B) GO cellular component analysis of DEPs related to cytoplasm-related terms in S-EVs. (C-D) GO biological process analysis and KEGG enrichment analysis of the DEPs in S-EVs (vs. L-EVs) related to muscle components. (E-F) GO biological process analysis and KEGG enrichment analysis of the DEPs in S-EVs (vs. L-EVs) related to cytoskeleton components. (G-H) GO biological process analysis and KEGG enrichment analysis of the DEPs in S-EVs (vs. L-EVs) related to ER components.

In addition, ER-related components were among the major enriched terms in S-EVs compared with L-EVs (Figure 3A). The ER plays an essential role in mediating cellular proliferation and survival by regulating protein processing and folding [24, 25]. GO analysis revealed that the S-EV proteins associated with ER-related components were involved in biological processes related to protein transport, modification and folding, and protein insertion and signaling (Figure 3G). KEGG analysis revealed that these S-EV proteins were involved in pathways related to motor proteins, the regulation of actin cytoskeleton, tight junction, adherens junction, and focal adhesion (Figure 3H), which again suggested that S-EVs might engage mainly in muscle function regulation. However, unlike our tissue S-EV results, a previous study reported that L-EVs isolated from dendritic cells (pelleted at 16,500 ×g) are more strongly associated with endoplasmic reticulum (ER) proteins than S-EVs [15]. This effect may be due to differences in EV origins (cells vs. tissues) and/or isolation methods, which suggests high heterogeneity of tissue-derived EVs. In addition, a few proteins upregulated in S-EVs (vs. L-EVs) were involved in certain metabolic pathways and signal transduction (Figure 3C and 3F). This effect may be because S-EVs and L-EVs might share some similar compositions or the potential overlap of some EV populations between the L-EV fraction and the S-EV fraction via the UC-based separation method (Figure 1H and Figure S1). Together, these results suggest that S-EVs carry abundant original muscle cell-specific compositions and might profoundly engage in regulating muscle development and function.

### L-EVs and S-EVs have distinct membrane proteins and cellular uptake patterns

In addition to their luminal cargos, EVs also carry many surface proteins that mediate interactions between EVs and target cells, influencing their cellular uptake, tissue selection/retention, and biological effects [26, 27]. For example, EVs from senescent fibroblasts are inefficiently internalized by fibroblasts and HeLa cells due to their enriched surface DPP4 protein [27]. The ability of breast cancer cell-derived EVs to promote cancer metastasis is also affected by surface integrins (e.g., ITGB3) [28]. Thus, the surface protein profiles of L-EVs and S-EVs were analyzed via a public surfaceome database [29] (Figure 4A). A Venn diagram revealed 302 surface proteins on the surface of muscle tissue-derived EVs (Figure 4B), of which 20 proteins were upregulated on L-EVs (vs. S-EVs) and 54 proteins were upregulated on S-EVs (vs. L-EVs; Figure 4C). Clusters of cell differentiation (CD) proteins (belonging to the tetraspanin family) and GPI-anchored proteins (GPI-APs) constitute a unique subset of EV-associated proteins, and they have been recognized as bona fide surface proteins [30]. Compared with L-EVs, S-EVs expressed higher levels of CD proteins (CD80, Tfrc, Ptgfrn, etc.) and GPI-APs (Hyal2, Ca4, CD48, etc.) (Figure 4C-4D, Figure S3A-3B). The elevated surface proteins of S-EVs (vs. L-EVs) were enriched in pathways associated with GPI anchor biosynthesis, attachment of the GPI anchor to uPAR, protein digestion and absorption, and trans-Golgi network vesicle budding (Figure 4E and 4F), whereas the elevated surface proteins of L-EVs (vs. S-EVs) were enriched in pathways related to oxidative phosphorylation, cholesterol metabolism, and the mitochondrial transport process (Figure 4G and 4H). CD protein (e.g., CD151 and Tspan8)-mediated sorting of adhesion receptors to EVs promotes the targeting and binding of pancreatic cancer cell-derived EVs to recipient cells [31]. Thus, our findings suggest that S-EVs and L-EVs have distinct surface protein profiles, which might be associated with different cellular uptake action.

**Figure 4.**
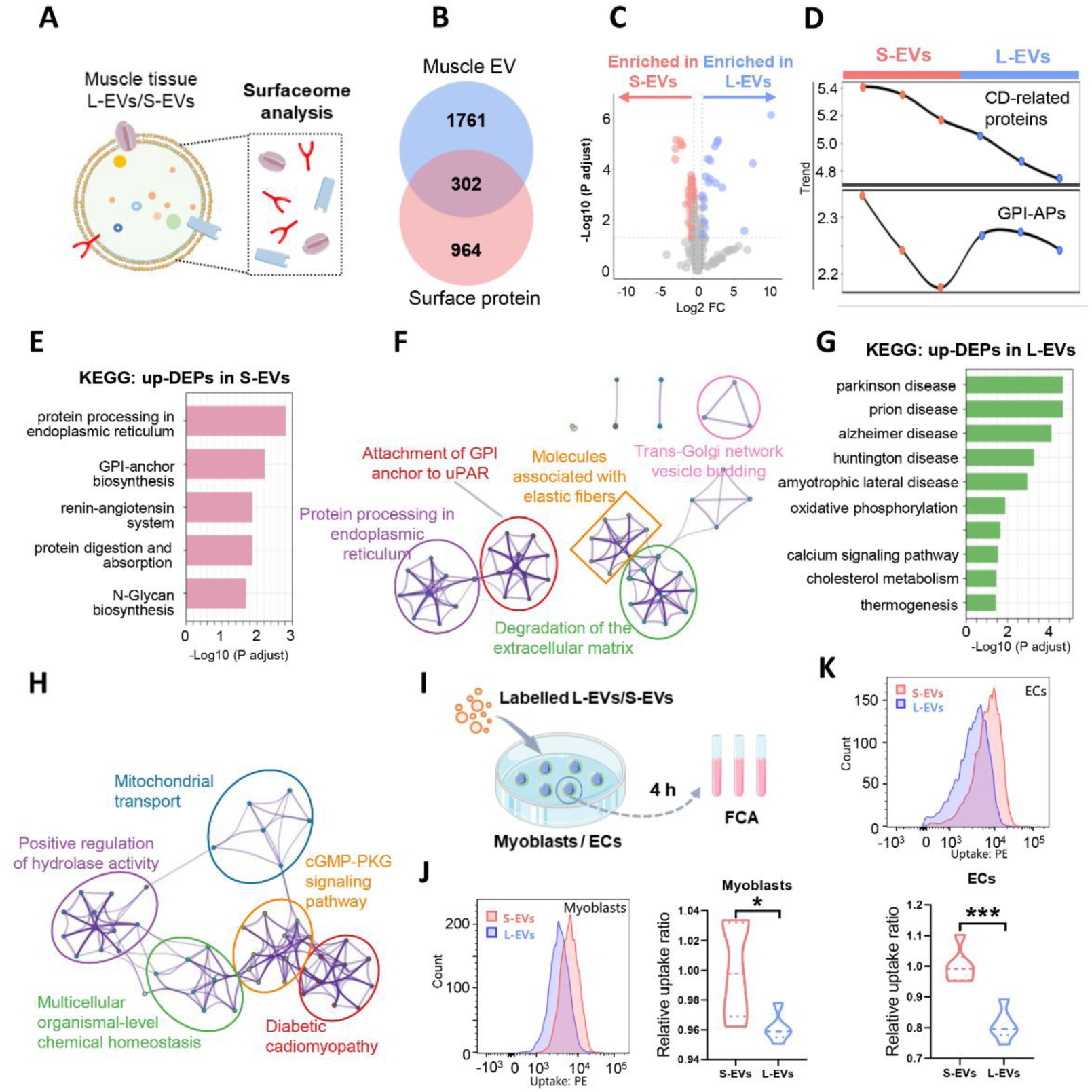
Evaluating surface protein profiles and cell uptake of tissue-EVs. (A) Schematic illustration of the strategies for analyzing EV surface proteins. (B) Venn diagram showing the surface proteins contained in muscle tissue-EVs (the overlap). (C) Volcano plot showing the surface DEPs between L-EVs and S-EVs (FC > 1.2 and p-adjusted < 0.05, n = 3). (D) Trend analysis of CD-related and GPI-AP expression levels between L-EVs and S-EVs. (E-F) KEGG enrichment and interaction network of the surface DEPs in S-EVs (vs. L-EVs). (G-H) KEGG enrichment and interaction network of the surface DEPs in L-EVs (vs. S-EVs). (I) Schematic illustration of the cellular uptake assay. (J-K) Evaluation of myoblast (J) and EC (K) uptake after incubation with Memglow-labeled EVs for 4 h via FCA. (n=3)

Skeletal muscle fibers mainly originate from myoblasts and the blood vascular network, which can support the development and normal function of muscle cells [32]. Additionally, we found that myoblasts and ECs might be the major origins of muscle tissue-EVs (Figure 1G). The binding and uptake of EVs by recipient cells is a critical step for their cell‒cell communication effect. The cellular uptake rates of tissue-EVs were assayed in mouse myoblasts and human ECs (Figure 4I). The Memglow (a membrane dye)-labeled S-EVs or L-EVs could be taken up by myoblasts or ECs, and the uptake rates of S-EVs were further higher than those of L-EVs in myoblasts or ECs (Figure 4J-K), suggesting that the enriched S-EV surface proteins, such as CD and GPI-APs, might facilitate their binding and uptake by recipient cells. Additionally, the uptake efficiency of L-EVs or S-EVs was greater in myoblasts (∼90% positive populations) than in ECs (∼60-70% positive populations) (Figure 4J-4K). It has been proposed that cell-derived EVs are prone to greater uptake by their parent cells [33], and this effect may be due to the muscle cell origin of tissue-EVs. These results indicate that different EV subpopulations have distinct surface proteins and cell uptake patterns and that harnessing this specific property might be promising for developing advanced targeted delivery vehicles for therapeutic purposes.

### L-EVs and S-EVs induce diverse biological responses in skeletal muscle cells

Having found that healthy muscle tissue-derived S-EVs and L-EVs have distinct compositions involved in different pathways, we next determined whether they induce diverse biological responses in recipient cells within the muscle tissue microenvironment. Myoblasts are a major type of skeletal muscle cell that can migrate to the region where myofibers form and differentiate into myotubes during muscle development [34]. The global responses of myoblasts to tissue-EVs were assessed via RNA-seq (Figure 5A). Notably, L-EV or S-EV treatment induced distinct gene expression patterns in myoblasts (Figure 5B). The DEGs induced by L-EVs were associated mainly with cell organelle components, such as the sarcoplasmic reticulum, receptor complex, and mitochondrial pyruvate dehydrogenase complex, whereas the DEGs induced by S-EV treatments were associated mainly with cellular structure components, such as nucleus, centrosome, and cytoplasmic vesicle (Figure S4A-S4D). Similarly, the DEGs of the L-EV group (vs. the S-EV group) were related primarily to organelles and the cytoplasm, whereas the DEGs of the S-EV group (vs. the L-EV group) were linked mainly to the nucleus and microtube-related structures (Figure 5C). Molecular function (MF) analysis revealed that the DEGs induced by L-EVs were involved primarily in metabolic regulation, such as fatty acid binding, glutathione activity, oxidoreductase activity, and NAD^+^ activity (Figure 5D), whereas the DEGs induced by S-EVs were related mainly to nucleic acid and cell structure processes, such as RNA polymerase II transcription factor activity, sequence-specific DNA binding, and tubulin binding (Figure 5E). These results suggest that L-EVs may function primarily in energy metabolism and signal transduction and that S-EVs might engage primarily in mediating cell proliferation or migration.

**Figure 5.**
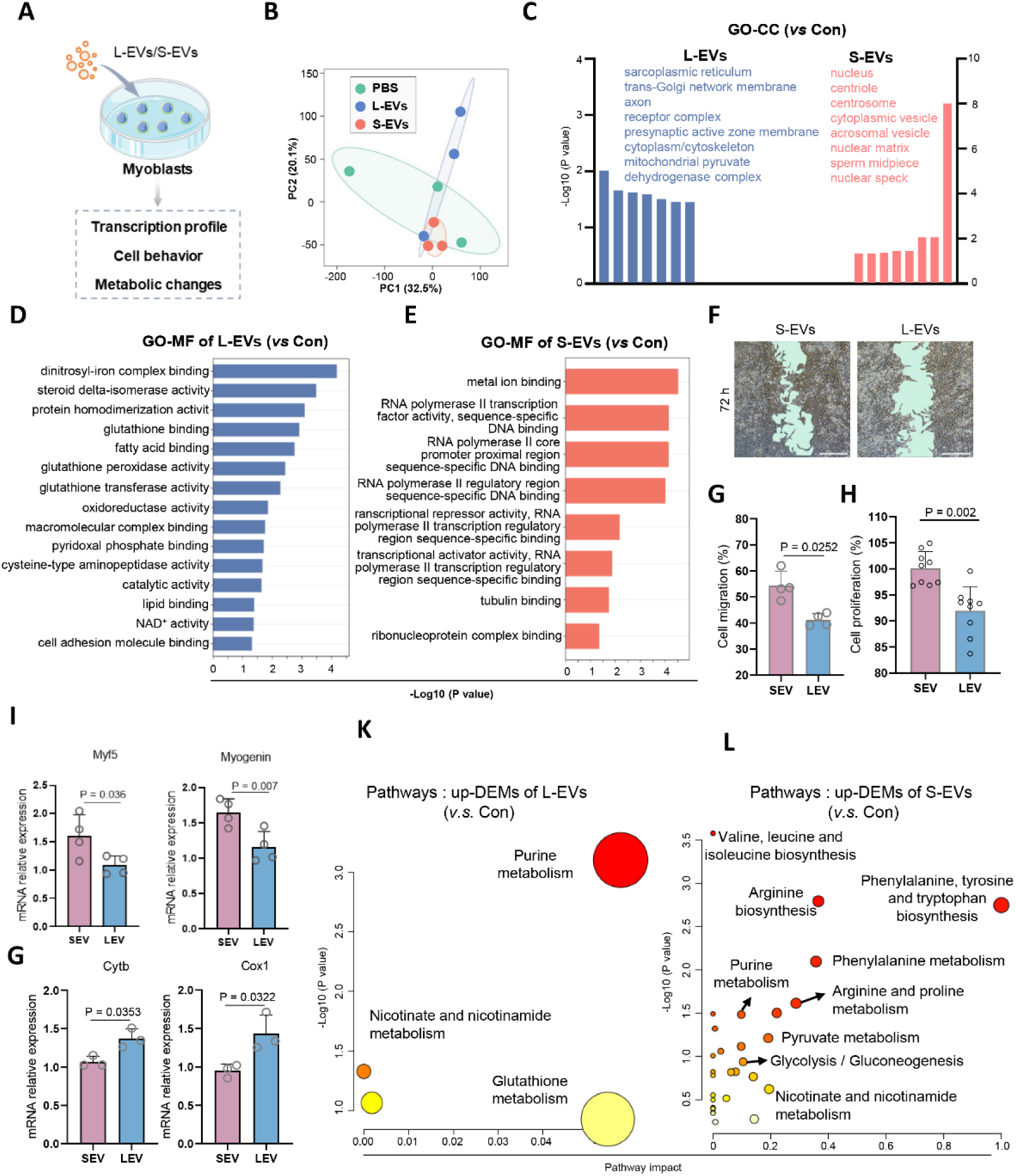
Evaluation tissue-EV-induced biological responses in myoblasts. (A) Schematic illustration of the functional experiments on myoblasts treated with or without EVs. (B) PCA scatter plot of different groups on the basis of RNA-seq data (n = 3). (C) GO cellular component analysis of the DEGs of myoblasts treated with L-EVs (left, vs. Con group) and S-EVs (right, vs. Con group). (D-E) GO molecular function analysis of the DEGs of myoblasts treated with L-EVs (D) and S-EVs (E). (F) Representative images of the migration of myoblasts treated with L-EVs or S-EVs (scale bar = 500 μm). (G) Quantification of the cell migration ratio in each group (n = 4). (H) Proliferation rate of myoblasts treated with L-EVs or S-EVs (n = 10). (I) qPCR analysis of the expression of genes related to differentiation (Myf5 and myogenin) in myoblasts treated with L-EVs or S-EVs. (G) qPCR analysis of mitochondrial gene expression (Cytb and Cox1) in myoblasts treated with L-EVs or S-EVs. (K-L) Pathway enrichment analysis of the DEMs enriched in myoblasts treated with L-EVs (K) and S-EVs (L).

Cell behaviors, such as the migration, proliferation, and differentiation of myoblasts, are vital events in the regulation of muscle development and homeostasis [34]. Thus, the impact of tissue-EVs on myoblast migration, proliferation or differentiation was evaluated *in vitro*. Myoblasts with S-EVs treatments showed higher levels of cell migration rates compared to L-EVs group (Figure 5F-G). Compared with L-EV treatment, S-EV treatment also led to higher cell proliferation rates in myoblasts (Figure 5H). It is well documented that Myf5 and myogenin are key regulators of myoblast differentiation and myogenic determination [35]. Our results revealed that the myoblasts in the S-EV group presented higher levels of Myf5 and myogenin expression than those in the L-EV group did (Figure 5I). Together with the proteomics results, these results suggest that S-EVs might function primarily in mediating cellular proliferation and differentiation. On the other hand, we found that the myoblasts in the L-EV group exhibited higher levels of mitochondria-related gene (e.g., Cytb and Cox1) expression than those in the S-EV group did (Figure 5G), suggesting a potential regulatory role of L-EVs in the metabolic metabolism of these cells.

Next, the global metabolic changes in myoblasts in response to tissue-EVs were analyzed via LC‒MS/MS-based targeted metabolomics. PCA plots and heatmaps revealed distinct metabolite profiles between the groups (Figure S5A-5B). Differentially expressed metabolites (DEMs) between the L-EV group and the S-EV group were identified (Figure S5C-5D). The DEMs induced by L-EVs were highly enriched in pathways related to energy and redox metabolism, particularly purine metabolism (Figure 5K). Purines constitute one of the major classes of metabolites that serve as the bases of energy units (e.g., ATP) and nucleotides to synthesize key components of cellular genetic materials [36], and purine metabolism is essential for controlling skeletal muscle functions (e.g., training status and performance) [37]. These results again suggest that L-EVs might engage in the metabolic dynamic regulation of muscle cells. S-EV treatment also moderately affected some pathways related to amino acid metabolism, but the pathway impact ratio was much lower than that in the L-EV group (Figure 5L). Together, L-EVs and S-EVs induce diverse biological responses in skeletal muscle cells, which may offer distinct potential for therapeutic development.

### L-EVs and S-EVs show functional heterogeneity in vascular endothelial cells

The development, function and injury repair processes of skeletal muscle are also dependent on blood vascular networks to provide necessary substances, such as oxygen and nutrients [32, 38]. We also found that ECs were one of the main cell sources of tissue-EVs in the muscle environment. Thus, the global impact of tissue-EVs on ECs was analyzed via RNA-seq, and each EV induced different gene expression profiles in ECs (Figure 6A-B). The upregulated DEGs in the L-EV group (vs. the Con group) were enriched in cellular components related to mitochondria (26.3%), nucleus (17.8%), and ribosome (9.5%) (Figure 6C). In line with the results from myoblasts, the DEGs were involved in mitochondrial processes and nucleic acid processing functions (Figure 6D-E). KEGG analysis revealed that the DEMs in the L-EV group (vs. the Con group) were involved mainly in pathways related to biomolecule synthesis and cellular functions, particularly purine metabolism (Figure S6A-S6B). We further verified the elevated expression levels of mitochondrial genes (e.g., Nd4l and Nd1) in ECs subjected to L-EV treatment (Figure 6H). Similarly, the mitochondria of ECs treated with L-EVs appeared to have enhanced fusion (elongated shape) compared with those in the control group (Figure 6I). These results collectively suggest that L-EVs induce changes in the mitochondrial dynamics of ECs, which might permit the mitochondria of ECs to maximize cellular energy production and adapt to increased cellular energy demands [39].

**Figure 6.**
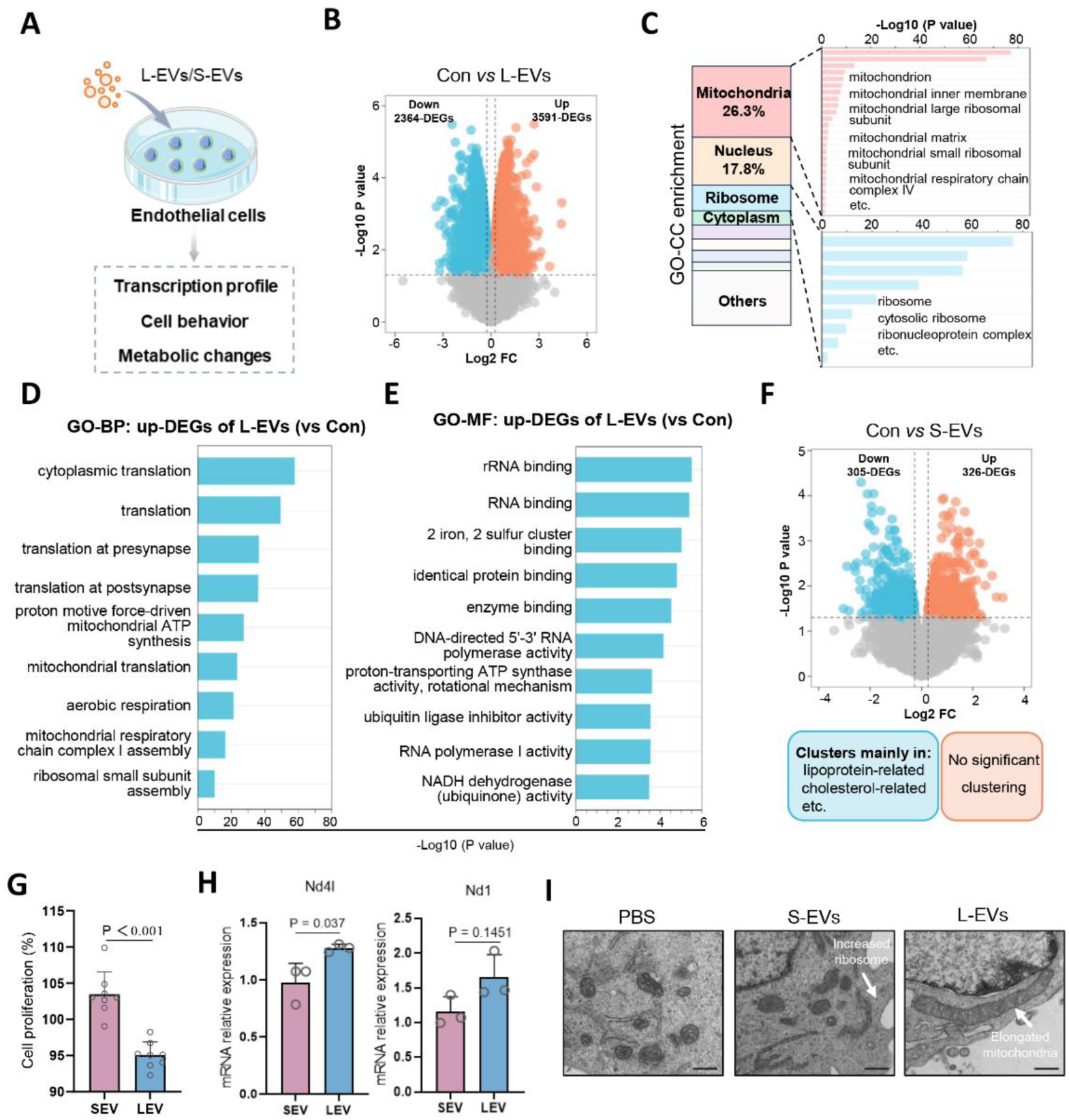
Evaluation of tissue-EV-induced biological responses in endothelial cells. (A) Schematic illustration of the functional experiments on ECs treated with/without EVs. (B) Volcano plot showing the DEGs in ECs treated with L-EVs (FC > 1.5 and p-value < 0.05, n = 3). (C) Sublocalization classification of GO-CC analysis of DEGs enriched in ECs treated with L-EVs (vs. Con). (D) GO biological process analysis of the DEGs in ECs treated with L-EVs (vs. the Con group). (E) GO molecular function analysis of the DEGs in ECs treated with L-EVs (vs. the Con group). (F) Volcano plot showing the DEGs of ECs treated with S-EVs (FC > 1.5 and p-value < 0.05, n = 3). (G) Proliferation rate of ECs treated with S-EVs or L-EVs (n = 8). (H) qPCR analysis of mitochondrial gene expression (Cytb and Cox1) in ECs treated with S-EVs or L-EVs (n = 3). (I) TEM images of ECs in each group (scale bar = 500 μm).

Although many DEGs (488 upregulated and 401 downregulated) were found in ECs treated with S-EVs compared with those in the control group (Figure 6F), no significantly enriched clusters were observed among the upregulated DEGs (Figure 6F). However, ECs treated with S-EVs presented higher cell proliferation rates than those in the L-EV group did (Figure 6G). In addition, similar to the results of S-EV treatment in myoblasts (Figure 5), S-EV treatment altered several metabolic pathways in ECs, but their pathway impact ratios were relatively low (Figure S6C). Ribosomes play crucial roles in mediating cell proliferation and differentiation since they are major sites of protein synthesis via the use of mRNAs and amino acids [40, 41]. We found that ECs of S-EVs group showed an increase trend in ribosome density compared to those of control group (Figure 6I), suggesting that the proliferative role of S-EVs is at least partly dependent on ribosome pathways. Together, these findings indicate that L-EVs and S-EVs play diverse roles in the regulation of cell metabolic states and phenotypes and that different tissue-EV subpopulations likely control tissue hemostasis in a coordinated manner. Owing to the intrinsic heterogeneity in the compositions and functions of tissue-EV subpopulations, a better understanding of their biological roles will provide insights into EV-mediated disease pathology and therapies.

Although remarkable progress has been made in the EV field, understandings on EV heterogeneity *in vivo* and the detailed role of EV subsets in the complicated tissue microenvironment remain challenging, which may hinder the further development of EV-based diagnostic and therapeutic applications. As one of the largest organs in the body, skeletal muscle tissue-derived EVs were selected as a proof-of-concept to investigate the compositional and functional heterogeneity of EV subpopulations *in vivo*. Importantly, different EV subpopulations (separated on the basis of only two size ranges) of healthy skeletal muscle tissues have shown distinct composition signatures, resulting in differences in the cellular uptake efficiency and biological responses of recipient cells (myoblasts and vascular ECs). In terms of future translation, S-EVs might be more suitable for biomarker development because of the enriched signatures from original cells, and L-EVs might be more ideal for therapeutic development for metabolic regulation because of their abundant mitochondrial contents. Moving forward, caution should be exercised when working with tissue-EVs, and such a reconsideration of EV subpopulation selection would aid in enhancing specificity and sensitivity when developing EV-based diagnostic modalities and reducing stochasticity when producing more tailored therapeutic EVs. In addition, evaluating the tissue selection and retention of different EV subpopulations *in vivo is worthwhile*. Nevertheless, this study highlights the compositional and functional heterogeneity of tissue EV subpopulations, and an in-depth understanding of this intrinsic property can promote the development of EV-based applications in the future.

## Materials and methods

### Animals

All animal experiments were performed following the Guidelines of the National Institutes of Health on the Use of Laboratory Animals and were approved by the Animal Care and Use Committee of West China Hospital, Sichuan University (Permit No. 20220210002). Healthy C57BL/6 mice (male, 6‒8 weeks) were purchased from Byrness Weil Biotech Ltd. (Chengdu, China) and housed in a special pathogen-free facility.

### Isolation of skeletal muscle tissue-EVs

Tissue-derived EVs were isolated from mouse skeletal muscle tissues according to previously reported methods with slight optimization[6]. In brief, skeletal muscle was dissociated from euthanized mice, cut into 1–3 mm^3^ pieces and washed with PBS. Then, the muscle pieces were digested with collagenase IV (1 mg/mL, 17104019, Gibco) and dispase (1 U/mL, D6430, Solarbio, Beijing, China) in a shaking incubator at 37°C for 2 h. After digestion, a large volume of precooled DMEM was used to stop the digestion, followed by centrifugation at 300 × g for 10 min to remove the tissue masses and 2,000 × g for 40 min at 4°C to remove the cells and other debris. The supernatant was subsequently ultracentrifuged at 30,000 × g for 60 min at 4°C in an SW32Ti rotor (Beckman Coulter, Brea, CA, USA) to obtain L-EVs, which were further ultracentrifuged at 110,000 × g for 70 min at 4°C to obtain S-EVs. The purified L-EV and S-EV preparations were resuspended in sterile PBS and stored at -80°C until further use. The protein concentrations of the EV samples were quantified via a BCA protein assay kit (CWBIO, Beijing, China). The particle concentrations of the EV samples were analyzed via a nanoparticle tracking analyzer (NTA, Particle Metrix, Meerbusch, Germany).

### Transmission electron microscope (TEM)

The micromorphology of muscle tissues, EVs or cells was observed via transmission electron microscopy (TEM) (HT7800, Hitachi, Ltd., Tokyo, Japan) as previously described [6, 42]. For the preparation of tissue TEM samples, muscle tissues were quickly fixed at 4°C in 3% glutaraldehyde and postfixed in 1% osmium tetroxide, followed by tissue dehydration and resin embedding. The embedded tissues were subsequently cut into ultrathin sections (70 nm), stained with 2% uranyl acetate and Reynolds lead citrate, and then observed via TEM. For observation of EV samples, the EV suspension was dropped on a carbon-coated copper grid, followed by washing with pure water to remove unattached EVs. The grid was then stained with 2% phosphotungstic acid and air-dried before being subjected to TEM. For observation of cell samples, cells treated with EVs (∼1×10^10^ particles/ml) for 24 h were collected, quickly fixed at 4°C in 3% glutaraldehyde and postfixed in 1% osmium tetroxide, followed by dehydration and embedding in resin. The embedded cells were subsequently cut into ultrathin sections (70 nm), stained with 2% uranyl acetate and Reynolds lead citrate, and then observed via TEM.

### LC‒MS/MS-based proteomic analysis of EVs

Total protein was extracted from EV samples via protein lysis buffer (8 M urea, 1% SDS) and quantified with a BCA kit (Thermo Fisher Scientific). EV protein samples (100 µg) were removed for reductive alkylation and digested with trypsin overnight at 37°C. The trypsin-digested peptides were desalted and drained with a vacuum concentrator. The peptides were subsequently analyzed via a VanquishNeo coupled with an Orbitrap Astral mass spectrometer (Thermo Fisher Scientific) at Majorbio Bio-Pharm Technology Co. Ltd. (Shanghai, China). Briefly, the peptides were dissolved in 0.1% formic acid and separated on an ES906 column (150 µm × 15 cm, Thermo Fisher Scientific) over a 60 SPD gradient at a flow rate of 500 nL/min. Data-independent acquisition (DIA) data were acquired via an Orbitrap Astral mass spectrometer, with MS data collected over an m/z range of 100-1700. Bioinformatic analysis of the proteomic data was performed with the Majorbio Cloud platform (https://cloud.majorbio.com). DEPs were identified using thresholds of fold change (FC) > 1.2 and p-adjusted < 0.05. PCA plots, volcano plots, and heatmaps were generated via a free online platform (https://www.omicsolution.org/wkomics/main/). GO analysis of the DEPs was performed via DAVID (https://david.ncifcrf.gov/) and STRING (https://cn.string-db.org/), with false discovery rate (FDR) < 0.05 considered significant. The surface data were generated via http://wlab.ethz.ch/surfaceome.

### Cell culture

Mouse myoblast cell lines (C2C12 cells) or human umbilical vein endothelial cell lines (human ECs, EA. hy926 cells) were cultured in DMEM (Gibco) supplemented with 10% FBS and 1% penicillin‒streptomycin. All the cells were cultured in an incubator under a humid atmosphere with 5% CO_2_ at 37°C.

### Cellular uptake assay of tissue-EVs

EVs were labeled with Memglow dye (2 μM; Cytoskeleton, Inc.) following the manufacturer’s instructions. The free dye was removed via ultracentrifugation. Myoblasts and ECs were incubated with dye-labeled EVs (20 μg/ml) or an equivalent amount of dye control for 4 h at 37°C with 5% CO_2_. After incubation, the cells were collected and washed with PBS. The positive signals of the stained cells were analyzed via a flow cytometer (CytoFLUX, Chengdu Legend-tech Biotech Co., ltd, China).

### RNA sequencing (RNA-seq) analysis

Mouse myoblast and human ECs were treated with PBS, L-EVs or S-EVs (∼1×10^10^ particles/ml) for 24 h. Then, the cells were collected for RNA-seq analysis. In brief, total RNA was extracted from the cells via TRIzol and then treated with DNase I (Takara, Shiga, Japan) to remove genomic DNA. The RNA-seq transcriptome library was constructed via a TruSeq™ RNA Sample Preparation Kit (Illumina, San Diego, CA, USA), and the paired-end RNA-seq sequencing library was sequenced on an Illumina HiSeq Xten/NovaSeq 6000 sequencer by Shanghai Majorbio Biopharm Technology Co., Ltd. (Shanghai, China). The DEGs with FC > 1.2 and p-value < 0.05 were considered significant. The PCA plot, volcano plot, and heatmap were generated via a free online platform (https://www.omicsolution.org/wkomics/main/). GO analysis of the DEGs was performed via DAVID (https://david.ncifcrf.gov/), and FDR < 0.05 was considered to indicate statistical significance.

### Cell migration assay

Mouse myoblasts were cultured in 12-well plates, and a straight line was scratched with a sterile 1 mL pipette tip when the cells reached 90% confluence. Then, the cells were treated with EVs (∼1×10^10^ particles/ml) and cultured in FBS-free DMEM for 72 h. Images of the narrow wound-like gaps were captured with an inverted microscope (Eclipse TS100, Nikon, Japan) and analyzed with ImageJ software (NIH, Bethesda, MD, USA).

### Cell proliferation assay

Mouse myoblast or human ECs were seeded into 96-well plates and treated with EVs (∼1×10^10^ particles/ml) for 24 h. Cell proliferation was measured via a CCK-8 Kit (Dojindo, Kumamoto, Japan). Briefly, CCK-8 solution was added to each well, and the plates were subsequently incubated for 40 min at 37°C. The absorbance at 450 nm was measured via a microplate reader (Tecan Group Ltd., Switzerland).

### Quantitative real-time polymerase chain reaction (qPCR)

Mouse myoblast or human ECs were seeded into 6-well plates and treated with EVs (∼1×10^10^ particles/ml) for 24 h. Total RNA was extracted from the cells via TRIzol reagent (Gibco), and cDNA was synthesized via a commercial kit (Vazyme, Nanjing, China). qPCR was performed with SYBR Green (Vazyme) on a CFX96 real-time PCR detection system (Bio-Rad, Hercules, CA, USA). The primers used in this study are listed in Table S1. The data were analyzed via Bio-Rad CFX Manager software, and the relative fold change in mRNA levels was determined via the delta‒delta Ct method with *Rps18* as the internal reference gene.

### LC‒MS/MS-based targeted metabolomics

Mouse myoblast or human ECs were seeded into 6-well plates and treated with EVs (∼5×10^9^ particle/ml) for 4 h. The intracellular metabolites of macrophages were extracted using methanol/H_2_O solution as previously described [42]. After washing with PBS, the cells in the wells of a 6-well plate were immediately extracted with buffer (1 mL, methanol/H_2_O = 8:2, v/v) and incubated at −20°C for 30 min. After centrifugation (13,000 rpm for 20 min at 4°C), 0.7 mL of the supernatant was collected and dried in a drier (Eppendorf, Fisher Scientific, Pittsburgh, PA) at 30°C for 4 h. The dried samples were reconstituted in 100 μL of hydrophilic interaction chromatography (HILIC) solution.

Targeted MS-based metabolomics was performed using an Agilent 1260 LC (Agilent Technologies, Santa Clara, CA) coupled with an AB Sciex Qtrap 5500 MS (AB Sciex, Toronto, Canada) system. The sample solution was injected into the LC‒MS/MS system and analyzed under positive (2 μl of sample solution) and negative (10 μl of sample solution) ion modes. The mobile phase, gradient conditions and MS parameters were set up as previously described[42]. Multiple reaction monitoring (MRM) mode was used to detect multiple metabolites. The metabolites were normalized to the cellular protein and FC > 1.2 and p-value < 0.05 were considered significant. PCA plots, and heatmaps were generated using a free online platform (https://www.omicsolution.org/wkomics/main/). The differentially expressed metabolites were analyzed and visualized using MetaboAnalyst (version 5.0, https://www.metaboanalyst.ca/MetaboAnalyst).

### Statistical analysis

All the data are presented as the means ± standard deviation (SD) and were analyzed using GraphPad software (version 8.0.2, IBM Corporation, USA) with t-tests (for two-group comparisons) or one-way analysis of variance (ANOVA) (for more than two group comparisons), and P < 0.05 was considered to indicate a significant difference. The data were obtained from at least three biological replicates, and “n” represents the number of independent samples for each group.

## Acknowledgements

This study was partly supported by the National Natural Science Foundation of China (32271438, 32071453, 82472129), Sichuan Science and Technology Program (2024NSFSC0586) and 1.3.5 Project for Disciplines of Excellence (ZYYC23001), West China Hospital of Sichuan University. The authors would like to thank Xijing Yang and Xiaoting Chen for the assistance in animal experiments, Ziwei Huang for assisting with the cellular experiments.

## Data and materials availability

All the data needed to evaluate the conclusions in the paper are presented in the paper and/or the Supplementary Materials.

