## Supplemental materials for "Unveiling the functional heterogeneity of endogenous tissue extracellular vesicles in skeletal muscle through multi-omics"

* Corresponding Author:

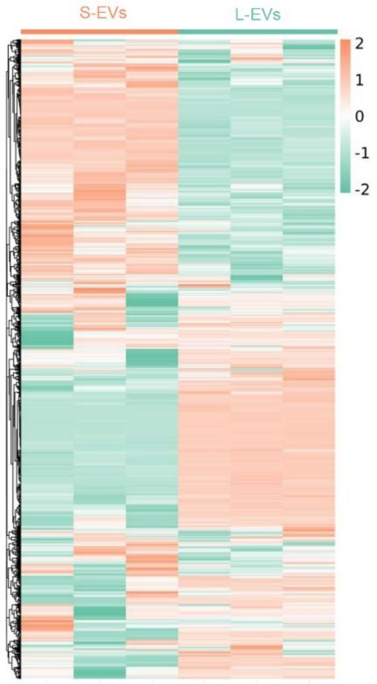


**Figure S1** Heatmap analysis of proteomic data from L-EVs and S-EVs from skeletal muscle tissues (n=3).

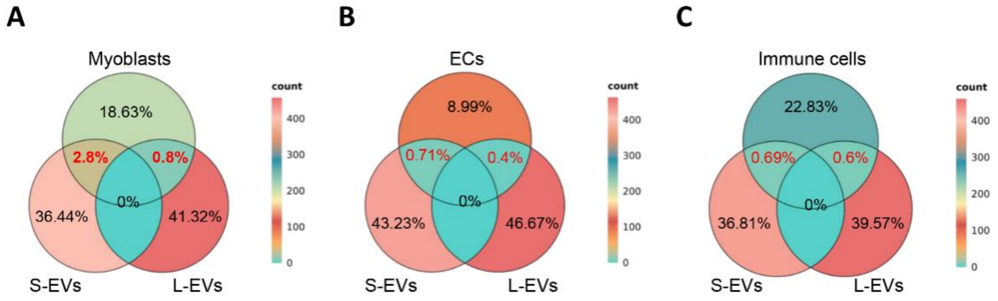


**Figure S2 Expression of the identified markers of the original cells in different EV subsets.** (A) Venn diagram showing the ratio of DEPs of S-EVs or L-EVs among the identified markers of myoblasts. (B) Venn diagram showing the ratio of DEPs of S-EVs or L-EVs among the identified markers of ECs. (C) Venn diagram showing the ratio of DEPs of S-EVs or L-EVs among the identified markers of immune cells.

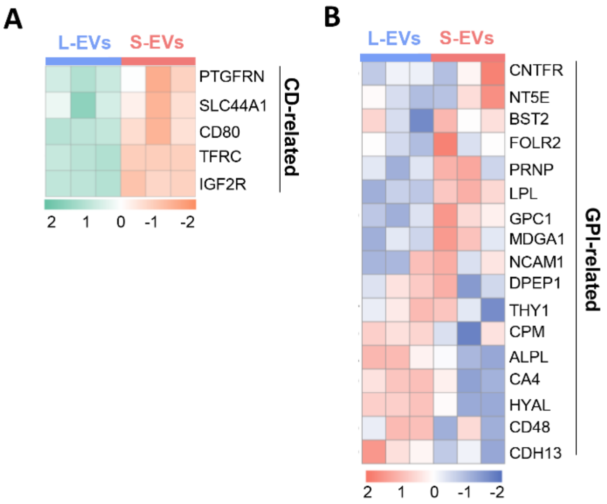


**Figure S3 Analysis of surface protein expression in tissue-derived EVs.** (A) Heatmap analysis of CD-related surface protein expression between L-EVs and S-EVs. (B) Heatmap analysis of GPI-related surface protein expression between L-EVs and S-EVs.

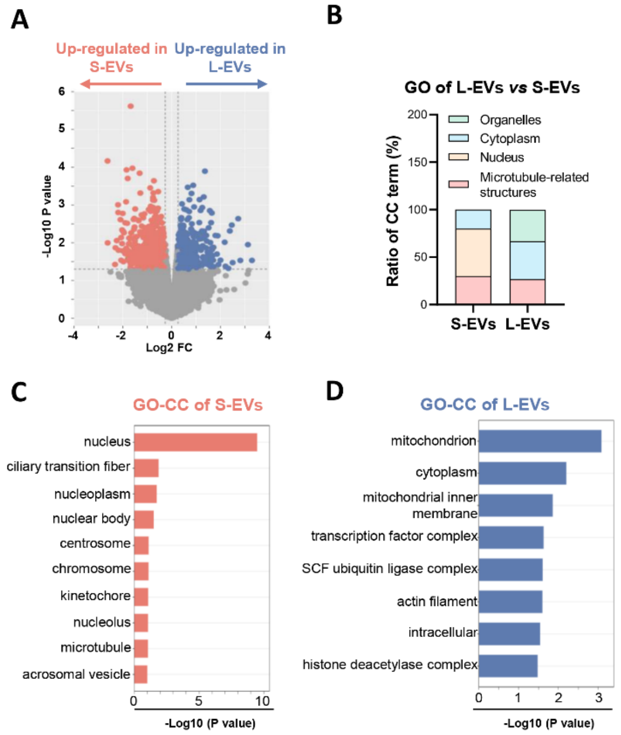


**Figure S4 Transcriptome analysis of myoblasts treated with tissue-derived EVs.**

(A) Volcano plot showing the DEGs in myoblasts between the L-EV- and S-EV-treated groups (FC > 1.5 and p-value < 0.05, n = 3). (B) Sublocalization classification of GO-CC analysis of DEGs in myoblasts treated with L-EVs and S-EVs. (C) GO cellular component analysis of DEGs in myoblasts treated with S-EVs (vs. L-EVs). (D) GO cellular component analysis of DEGs in myoblasts treated with L-EVs (vs. S-EVs).

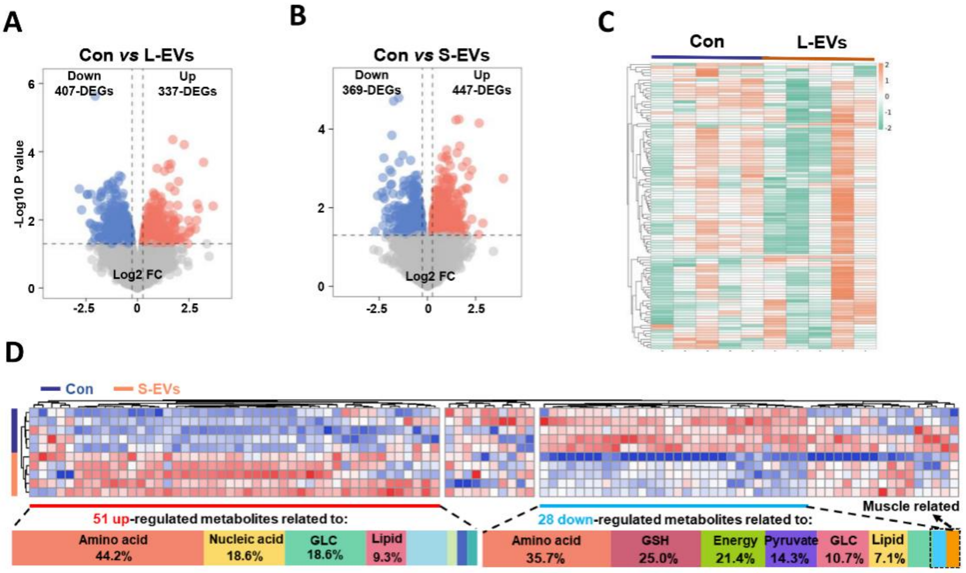


**Figure S5 Metabolic analysis of myoblasts treated with tissue-derived EVs.** (A) Volcano plot showing the DEMs in myoblasts treated with L-EVs (vs. Con, FC > 1.2 and p-value < 0.05, n = 5). (B) Volcano plot showing the DEMs in myoblasts treated with S-EVs (vs. Con, FC > 1.2 and p-value < 0.05, n = 5). (C) Heatmap analysis of DEMs in myoblasts treated with L-EVs (vs. Con). (D) Heatmap analysis of DEMs in myoblasts treated with S-EVs (vs. Con).

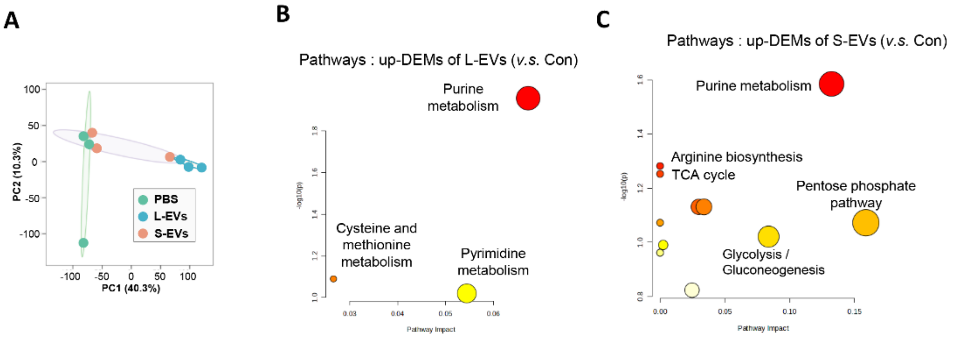


**Figure S6 Metabolic analysis of ECs treated with tissue-derived EVs.** (A) PCA scatter plot showing the differences in ECs between the different groups (n = 5). (B) Pathway enrichment analysis of the DEMs enriched in ECs treated with L-EVs (vs. Con). (C) Pathway enrichment analysis of the DEMs enriched in ECs treated with S-EVs (vs. Con).

**Table S1. The primer sequences used in this study.**

| Gene | Sequence 5'-3' | Species |
|---|---|---|
| Myf5 | Forward: ACCACCACCAACCCTAACCAGAG<br>Reverse: CGGCAGGCTGTAATAGTTCTCCAC | mouse |
| Myogein | Forward: GTCCCAACCCAGGAGATCATTTGC<br>Reverse: TCCTCCACCGTGATGCTGTCC | mouse |
| Cytb | Forward: CAAACCTCCTATCAGCCATCC<br>Reverse: AGCGAAGAATCGGGTCAAG | mouse |
| Cox1 | Forward: TACTATTCGGAGCCTGAGCG<br>Reverse: GTGTGATATGGTGGAGGGCA | mouse |
| RPS18 | Forward: TTCGCCATCACTGCCATTAAGGG<br>Reverse: ATCACTCGCTCCACCTCATCCTC | mouse |
| Nd4l | Forward: CCCACTCCCTCTTAGCCAATATT<br>Reverse: TAGGCCCACCGCTGCTT | human |
| ND1 | Forward: CCCATGGCCAACCTCCTACTCCTC<br>Reverse: AGCCCGTAGGGGCCTACAACG | human |
| Rps18 | Forward: GCAGAATCCACGCCAGTACAAG<br>Reverse: GCTTGTTGTCCAGACCATTGGC | human |
